# A third vaccination with a single T cell epitope protects against SARS-CoV-2 infection in the absence of neutralizing antibodies

**DOI:** 10.1101/2021.12.15.472838

**Authors:** Iris N. Pardieck, Esmé T.I. van der Gracht, Dominique M.B. Veerkamp, Felix M. Behr, Suzanne van Duikeren, Guillaume Beyrend, Jasper Rip, Reza Nadafi, Tetje C. van der Sluis, Elham Beyranvand Nejad, Nils Mülling, Dena J. Brasem, Marcel G.M. Camps, Sebenzile K. Myeni, Peter J. Bredenbeek, Marjolein Kikkert, Yeonsu Kim, Luka Cicin-Sain, Tamim Abdelaal, Klaas P.J.M. van Gisbergen, Kees L.M.C. Franken, Jan Wouter Drijfhout, Cornelius J.M. Melief, Gerben C.M. Zondag, Ferry Ossendorp, Ramon Arens

**Affiliations:** Department of Immunology, Leiden University Medical Center, Leiden, The Netherlands; Department of Medical Microbiology, Leiden University Medical Center, Leiden, The Netherlands; Department of Viral Immunology, Helmholtz Centre for Infection Research, Braunschweig, Germany; Delft Bioinformatics Lab, Delft University of Technology, Delft, The Netherlands; Department of Radiology, Leiden University Medical Center, Leiden, The Netherlands; Department of Hematopoiesis, Sanquin Research and Landsteiner Laboratory, Amsterdam, The Netherlands; Immunetune BV, Leiden, The Netherlands; ISA Pharmaceuticals BV, Leiden, The Netherlands

## Abstract

Understanding the mechanisms and impact of booster vaccinations can facilitate decisions on vaccination programmes. This study shows that three doses of the same synthetic peptide vaccine eliciting an exclusive CD8^+^ T cell response against one SARS-CoV-2 Spike epitope protected all mice against lethal SARS-CoV-2 infection in the K18-hACE2 transgenic mouse model in the absence of neutralizing antibodies, while only a second vaccination with this T cell vaccine was insufficient to provide protection. The third vaccine dose of the single T cell epitope peptide resulted in superior generation of effector-memory T cells in the circulation and tissue-resident memory T (T_RM_) cells, and these tertiary vaccine-specific CD8^+^ T cells were characterized by enhanced polyfunctional cytokine production. Moreover, fate mapping showed that a substantial fraction of the tertiary effector-memory CD8^+^ T cells developed from remigrated T_RM_ cells. Thus, repeated booster vaccinations quantitatively and qualitatively improve the CD8^+^ T cell response leading to protection against otherwise lethal SARS-CoV-2 infection.

**Summary:** A third dose with a single T cell epitope-vaccine promotes a strong increase in tissue-resident memory CD8^+^ T cells and fully protects against SARS-CoV-2 infection, while single B cell epitope-eliciting vaccines are unable to provide protection.

## Introduction

The coronavirus disease 2019 (COVID-19) pandemic, caused by the β-coronavirus severe acute respiratory syndrome coronavirus 2 (SARS-CoV-2), remains a global health emergency. Vaccines eliciting neutralizing antibodies against the Spike protein of SARS-CoV-2 have shown high effectiveness (1, 2). However, the level of neutralizing antibodies declines after vaccination (3) and at-risk patient groups, including transplant recipients, individuals suffering from B cell leukemia, and auto-immune disease patients on selected immunosuppressive regimens (such as B cell depleting reagents), exhibit lower humoral and cellular immunity after vaccination (4–6). Moreover, the mutation rate of coronaviruses is substantial and certain mutations in the Spike protein of SARS-CoV-2 have emerged that can lead to a decline in neutralizing antibody-mediated protection (7, 8). In order to address this clinical problem, a third vaccination for solid organ transplant recipients is becoming the standard of care in more and more countries, and supplementary T cell-focused strategies have been suggested (9, 10).

While boosters with the current vaccines likely lead to the generation of enhanced levels of neutralizing antibodies, it is uncertain if this will improve protection in the aforementioned risk groups and provide protection against the emergence of novel SARS-related viruses. However, a third vaccination may also lead to enhanced T cell responses, which is expected to contribute to the control of SARS-CoV-2 infection as evidenced by their association to disease severity and protection in humans (11–16) and animal models (17, 18). Moreover, T cell-focused vaccines can be directed to more conserved regions of the coronavirus, which may lead to a broad T cell-based cross-protection against multiple coronavirus strains and likely independently expands the protection provided by antibodies. Such T cell-mediated protection is important given the potential emergence of similar viruses in the future that pose a significant threat to global public health. Previous studies showed that T cells can mediate protection by themselves against SARS-CoV-1 (19, 20) but whether T cell-eliciting vaccines can protect against SARS-CoV-2 is unclear. Here, we demonstrate that a peptide vaccine exclusively eliciting CD8^+^ T cell responses against a single epitope present in the Spike protein delivers full protection against SARS-CoV-2, provided that the vaccine was administered in a double booster regimen, leading to the induction of high numbers of circulating and tissue-resident memory T (T_RM_) cells. A third vaccination was especially critical to achieve cytokine polyfunctionality and induction of elevated T_RM_ cell numbers in the lungs and liver, and additionally fueled retrograde migration of T_RM_ cells into the circulation. Thus, booster vaccinations eliciting strong CD8^+^ T cell responses are a promising strategy against coronavirus-mediated disease independent of neutralizing antibodies.

## Results

### Linear B cell epitope vaccines are not effective against SARS-CoV-2 infection

Single B and T cell epitope vaccines can provide effective approaches against a variety of pathogens and malignancies (21–24). To investigate the efficacy of linear B cell epitope vaccines against SARS-CoV-2 infection, we selected five different linear epitopes present in the Spike protein that were considered to potentially elicit protective antibodies (25–27). To test their immunogenicity, C57BL/6 mice were vaccinated with synthetic long peptide (SLP)-based vaccines each containing a single B cell epitope. After three vaccinations, however, these SLP vaccines did not elicit Spike-specific antibody responses (**Supplementary Fig 1A**). To improve this synthetic vaccine, we hypothesized that lack of CD4^+^ T cell help may underlie the ineffective antibody responses. In line with that, addition of SLPs containing the universal helper T-cell epitope PADRE or coupling of the PADRE epitope to the linear B cell epitopes, improved the Spike-specific IgG antibody response (**Fig 1A, Supplementary Fig 1B**). Next, we tested the particular influence of the adjuvants CpG and Incomplete Freund’s Adjuvant (IFA) on the specific antibody induction of the B cell-SLP vaccines. The combination of these adjuvants acted synergistically to elicit high levels of antibodies against a single epitope present in the Spike protein, which correlated with a higher induction of PADRE-specific CD4^+^ T cell responses (**Fig 1B-D**). Depletion of CD4^+^ T cells during the vaccination period resulted in a decreased spike-specific antibody response (**Supplementary Fig 1C**), highlighting the need for CD4^+^ T cells for an optimal IgG response. CD8^+^ T cell responses were not induced by the B cell-SLP vaccines (**Supplementary Fig 1D**).

**Figure 1.**
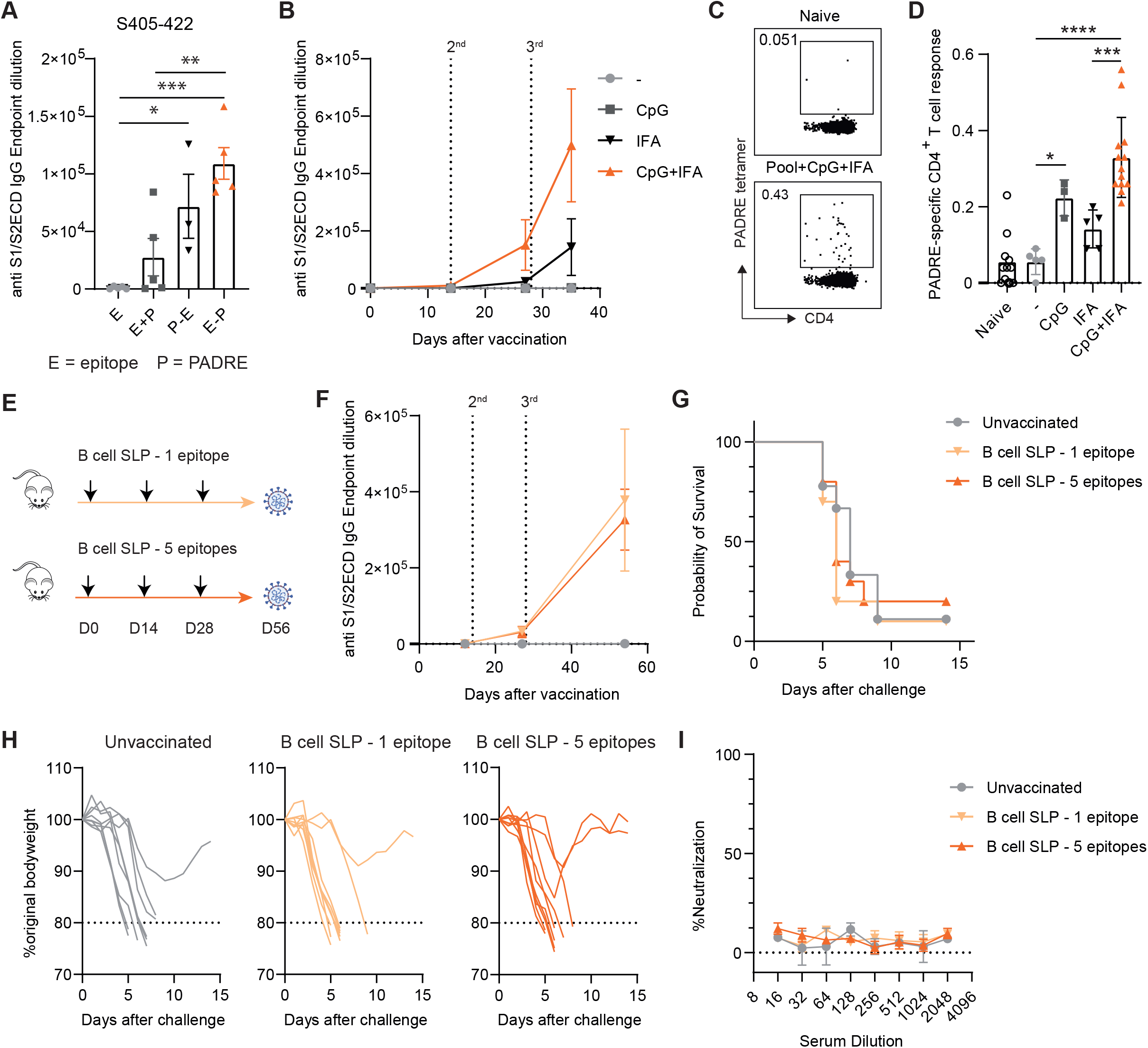
Linear B cell epitope vaccines are not effective against SARS-CoV-2 infection. **(A)** C57BL/6 mice were vaccinated subcutaneously (s.c.) on day 0 (1^st^ vaccination), day 14 (2^nd^ vaccination) and day 28 (3^rd^ vaccination) with synthetic long peptides (SLPs) consisting of PADRE-coupled linear B cell epitopes (E-P) or equimolar SLP controls containing the epitope alone (E), or the epitope combined with PADRE (E + P) adjuvanted with CpG and Inactivated Freund’s Adjuvant (IFA). Endpoint dilutions of Spike (S1/S2 ectodomain)-specific IgG antibodies in blood at day 42 after vaccination with Spike_405-422_-SLP are shown. Bar graphs represent means ± SEM. Symbols represent individual mice. One-way ANOVA was performed to determine statistical significance. **(B)** Spike S1/S2 ectodomain-specific IgG antibody kinetics in blood at indicated days after vaccination. Data is represented as mean ± SEM. **(C)** Representative flow cytometry plots of PADRE-specific CD4^+^ T cells determined by MHC class II tetramer staining on day 21 after the 1^st^ vaccination with SLPs consisting of PADRE-coupled linear B cell epitopes adjuvanted or not with CpG and/or Inactivated Freund’s Adjuvant (IFA). **(D)** PADRE-specific CD4^+^ T cell response on day 21 after vaccination as described in (C). Bar graphs represent means ± SEM. Symbols represent individual mice. One-way ANOVA was performed to determine statistical significance. **(E-I)** K18-hACE2 transgenic mice were vaccinated on day 0 (1^st^ vaccination), day 14 (2^nd^ vaccination) and day 28 (3^rd^ vaccination) with one or five SLP vaccines containing a PADRE-coupled linear B cell epitope adjuvanted with CpG and IFA. Four weeks after the 3^rd^ vaccination, mice were intranasally infected with 5000 PFU of SARS-CoV-2. **(F)** Spike S1/S2 domain-specific IgG antibody kinetics in blood at indicated days after B cell-SLP vaccination. Data is represented as mean ± SEM. **(G)** Survival graph of SARS-CoV-2 challenged K18-hACE2 transgenic mice after B cell-SLP vaccination. Significance was determined by a log-rank test and corrected for multiple comparisons. **(H)** Weight loss kinetics in time of SARS-CoV-2 challenged K18-hACE2 transgenic mice after B cell-SLP vaccination. **(I)** SARS-CoV-2 neutralizing capacity by antibodies after B cell-SLP vaccination on day 42 was measured by WT VNA determining the inhibition of the cytopathic effect of a SARS-CoV-2 isolate on Vero-E6 cells. *P<0.05, **P<0.01, ***P<0.001, ****P<0.0001.

To determine whether immunization with optimized linear B cell epitope vaccines enables protective immune responses against SARS-CoV-2, we used transgenic mice expressing the human angiotensin I-converting enzyme 2 (ACE2) receptor under the regulation of the cytokeratin 18 (K18) promoter, which develop severe lung disease in response to SARS-CoV-2 infection (28). The K18-hACE2 mice were vaccinated three times in a two week interval with either one CpG/IFA adjuvanted PADRE-coupled B cell-SLP vaccine or a combination of the five different B cell-SLP vaccines (**Fig 1E**). Spike-specific IgG antibody responses were elicited and reached high titers after the third (3^rd^) vaccination (**Fig 1F**). Four weeks after the 3^rd^ vaccination, the K18-hACE2 mice were challenged intranasally with a lethal dose of SARS-CoV-2 and body weight was measured daily as a parameter of disease. Both unvaccinated mice and mice that were vaccinated three times with the B cell-SLPs succumbed to SARS-CoV-2 infection (**Fig 1G, 1H**), showing that SLP vaccination with these linear B cell epitopes was not able to protect against SARS-CoV-2 infection. This inability of protection correlated with the deficiency of antibodies against the single epitopes to neutralize virus cell entry (**Fig 1I**). In contrast, a DNA vaccine encoding the entire Spike protein fully protected K18-hACE2 transgenic mice and this correlated with the ability to elicit neutralizing Spike-specific antibodies (**Supplementary Fig 1E**-**1H**). In addition, this DNA vaccine also induced a Spike-specific CD4^+^ and CD8^+^ T cell response (**Supplementary Fig 1I**). Overall, these results indicate that antibody responses to these single linear B cell epitopes are inferior for induction of neutralizing antibodies, although, we cannot exclude the possibility that single linear B cell epitopes exist that can elicit neutralizing antibodies. Nevertheless, antibody responses induced by vaccine platforms encoding properly folded proteins presenting conformational B cell epitopes are able to generate neutralizing antibodies and provide protection against SARS-CoV-2 infection.

### A third vaccination with a peptide harboring a single T cell epitope protects against SARS-CoV-2 challenge

To determine whether an immune response induced by a single CD8^+^ T cell epitope vaccine was able to protect against SARS-CoV-2 infection, K18-hACE2 transgenic mice were vaccinated once, twice or three times with an SLP encoding an immunodominant CD8^+^ T cell epitope present in the Spike protein of SARS-CoV-1 and SARS-CoV-2 (29). Five weeks after the final vaccination, mice were intranasally challenged with a lethal dose of SARS-CoV-2 (**Fig 2A**). SLP vaccination induced a Spike_539-546_-specific CD8^+^ T cell response in all mice, which increased with each booster immunization and remained elevated after contraction (**Fig 2B-D**). CD4^+^ T cell responses and spike-specific antibodies were not induced (**Supplementary Fig 1J, 1K**). Whereas 75% of the non-vaccinated mice succumbed to SARS-CoV-2 challenge, 66% and 50% of the mice that received one or two vaccinations, respectively, succumbed. Strikingly, all mice that received a 3^rd^ vaccine dose recovered from the SARS-CoV-2 challenge despite initial weight loss (**Fig 2E, 2F**). Thus, a third vaccination with an SLP containing a single CD8^+^ T cell epitope protects against lethal SARS-CoV-2 infection.

**Figure 2.**
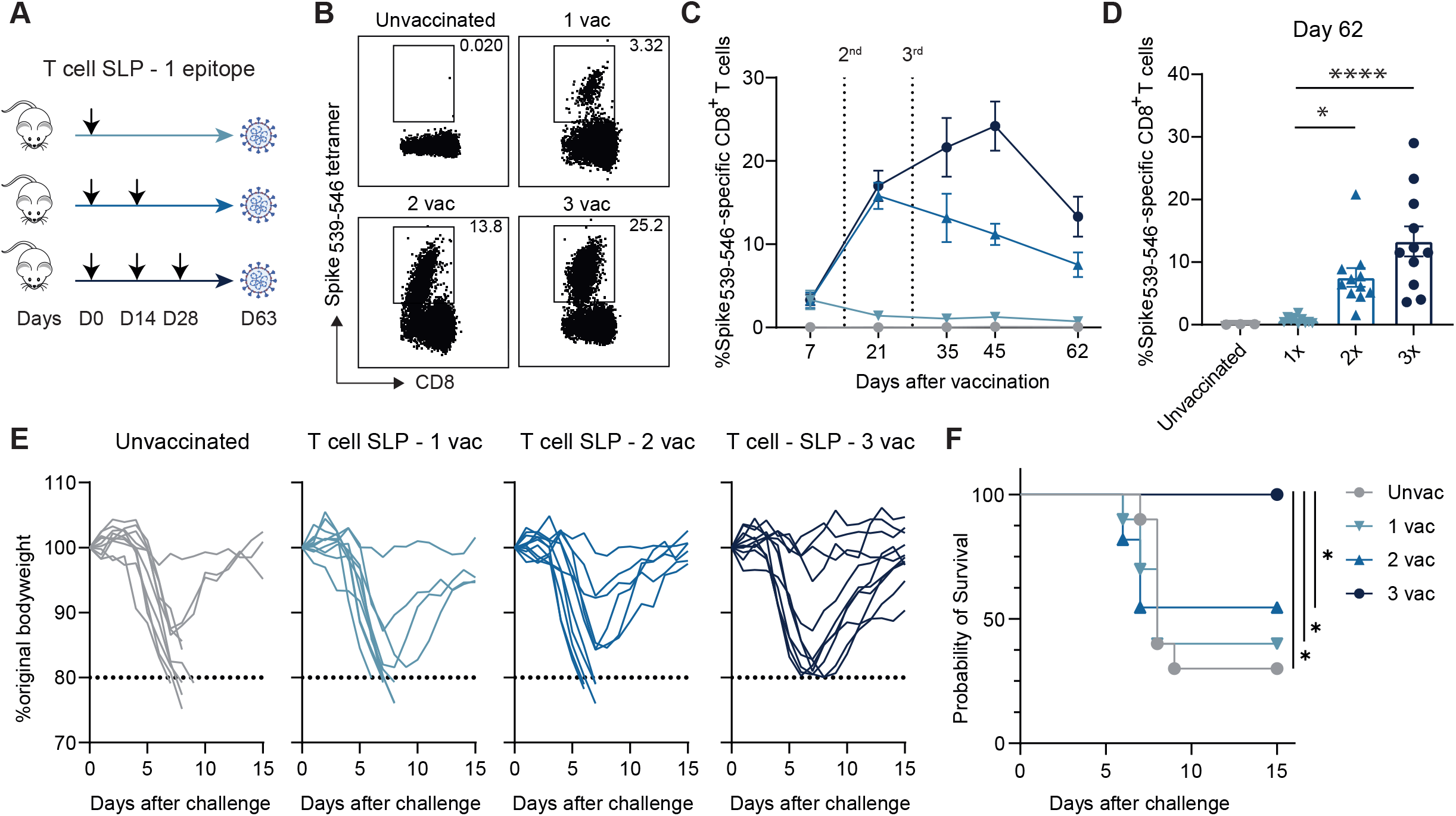
A third vaccination with a single T cell epitope protects against SARS-CoV-2 challenge. **(A)** K18-hACE2 transgenic mice were vaccinated subcutaneously on day 0 (1^st^ vaccination), day 14 (2^nd^ vaccination) and day 28 (3^rd^ vaccination) with the Spike_539-546_-SLP vaccine adjuvanted with CpG. The Spike_539-546_-specific CD8^+^ T cell response was determined in blood in time. Five weeks after the final vaccination, mice were intranasally infected with 5000 PFU of SARS-CoV-2, and monitored for weight loss and signs of clinical discomfort. **(B)** Representative flow cytometry plots of Spike_539-546_-specific CD8^+^ T cells determined in the blood circulation at day 45 by MHC class I tetramer staining. **(C)** Spike_539-546_-specific CD8^+^ T cell kinetics in blood at indicated days after vaccination. Data is represented as mean ± SEM. **(D)** Spike_539-546_-specific CD8^+^ T cells in blood at day 62 after vaccination. Bar graphs represent means ± SEM. Symbols represent individual mice. One-way ANOVA was performed to determine statistical significance. **(E)** Weight loss kinetics in time of SARS-CoV-2 challenged K18-hACE2 transgenic mice after T cell epitope SLP vaccination. **(F)** Survival graph of SARS-CoV-2 challenged K18-hACE2 transgenic mice after T cell SLP vaccination. Significance was determined by a log-rank test and corrected for multiple comparisons. *P<0.05, ****P<0.0001.

### Enhanced circulating and tissue-resident memory CD8^+^ T cell formation upon a third vaccination

Next, we aimed to determine the impact and underlying mechanisms of sequential vaccinations on the memory differentiation of the elicited antigen-specific CD8^+^ T cells. Three times vaccinated mice had higher frequencies of Spike_539-546_-specific CD8^+^ T cells in the spleen compared to two times vaccinated mice, which in turn had higher frequencies compared to once vaccinated mice (**Fig 3A**). Remarkably, in the liver and lungs, the 3^rd^ vaccination resulted also in a notably higher Spike_539-546_-specific CD8^+^ T cell response compared to the single or double vaccination. In the liver, this difference was mainly due to an increment of the CD69^+^ T_RM_ cells and in the lungs due to enhancement of both the circulating and T_RM_ cells (**Fig 3A, Supplementary Fig 2A**). To investigate the effector potential of these Spike_539-546_-specific CD8^+^ T cells, the polyfunctional cytokine production capacity of the vaccine-specific CD8^+^ T cells was determined. The 2^nd^ vaccination markedly increased the magnitude of the polycytokine producing CD8^+^ T cells (IFN-γ^+^/TNF^+^, IFN-γ^+^/TNF^+^/IL-2^+^) in spleen, liver and lungs, which was further increased substantially by the 3^rd^ vaccination. (**Fig 3B**).

**Figure 3.**
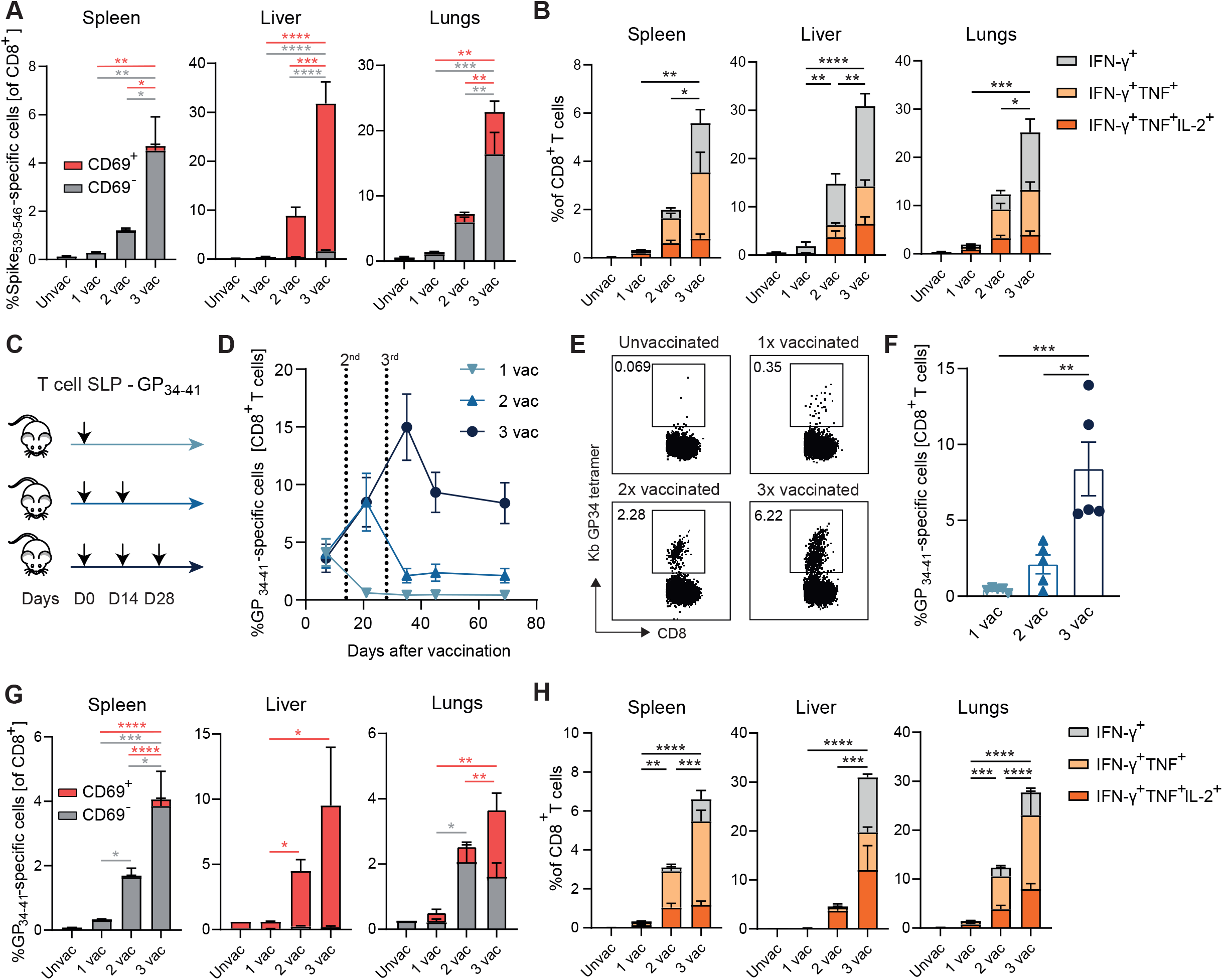
Enhanced effector-memory and tissue-resident memory CD8^+^ T cell formation upon a third vaccination. **(A)** Frequencies of CD69^+^ and CD69^-^ Spike_539-546_-specific CD8^+^ T cells in the CD8^+^ T cell population in the spleen, liver and lungs at day 70 after 1, 2 or 3 vaccinations. Bar graphs represent means ± SEM. One-way ANOVA was performed to determine statistical significance. **(B)** Intracellular cytokine production of CD8^+^ T cells upon stimulation with the Spike_539-546_ peptide epitope in different tissues on day 70 after SLP vaccination. Data is represented as mean ± SEM. One-way ANOVA was performed to determine statistical significance. **(C)** C57BL/6 mice were vaccinated with the GP_34-41_-SLP vaccine adjuvanted with CpG in a prime-boost-boost regimen with 2 week intervals. **(D)** GP_34-41_-specific CD8^+^ T cell kinetics in blood at indicated days after vaccination. Data is represented as mean ± SEM. **(E)** Representative flow cytometry plots of GP_34-4_ι-specific CD8^+^ T cells determined by MHC class I tetramer staining. **(F)** GP_34-41_-specific CD8^+^ T cells in blood at day 69 after vaccination. Bar graphs represent means ± SEM. Symbols represent individual mice. One-way ANOVA was performed to determine statistical significance. **(G)** Frequencies of CD69^+^ and CD69^-^ GP_34-41_-specific CD8^+^ T cells in the CD8^+^ T cell population in the spleen, liver and lungs at day 66 after 1, 2 or 3 vaccinations. Bar graphs represent mean ± SEM. One-way ANOVA was performed to determine statistical significance. **(H)** Intracellular cytokine production of CD8^+^ T cells upon stimulation with the GP_34-41_ peptide epitope in different tissues on day 66 after SLP vaccination. Data is represented as mean ± SEM. One-way ANOVA was performed to determine statistical significance. *P<0.05, **P<0.01, ***P<0.001, ****P<0.0001.

To validate these findings for other viral epitopes, we vaccinated mice with SLP vaccines containing the model antigen GP_34-41_ derived from lymphocytic choriomeningitis virus (LCMV) (**Fig 3C**). As observed with the Spike_539-546_-SLP immunization, each subsequent vaccination with GP_34-41_-SLP, increased the magnitude of the vaccine-specific CD8^+^ T cell response, and this response remained higher after contraction (**Fig 3D-F**). Moreover, the circulating GP_34-41_-specific CD8^+^ T cells in the spleen where significantly increased upon each vaccination, while in the liver and lungs, the 3^rd^ vaccination resulted in a much higher magnitude of the GP_34-41_-specific CD8^+^ T_RM_ cells (**Fig 3G, Supplementary Fig 2B**). In addition, these increments were also reflected in increased polyfunctional cytokine producing GP_34-41_-specific CD8^+^ T cells in the spleen, liver and lungs (**Fig 3H**). Thus, sequential booster vaccinations result in programming significant and durable expansion of polyfunctional vaccine-specific CD8^+^ T cells.

### Boosting impact on the phenotype of vaccine-specific CD8^+^ T cells

Next, we aimed to examine the phenotype of the vaccine-boosted CD8^+^ T cells. We first performed analysis of these cells in the blood circulation using the canonical markers CD62L, CD44 and KLRG1, which define their state of differentiation. Both conventional flow cytometric analysis and analysis using *Cytosplore*, which incorporates approximated t-distributed stochastic neighbor embedding (A-tSNE) algorithms for subset definition (30), highlighted that the memory phenotype of the GP_34-41_-specific CD8^+^ T cells in blood is clearly skewed towards an effector-memory like state (CD44^high^CD62L^-^KLRG1^+^) by one booster vaccination, and that the 2^nd^ booster vaccination skewed this phenotype slightly further (**Fig 4A, 4B**).

**Figure 4.**
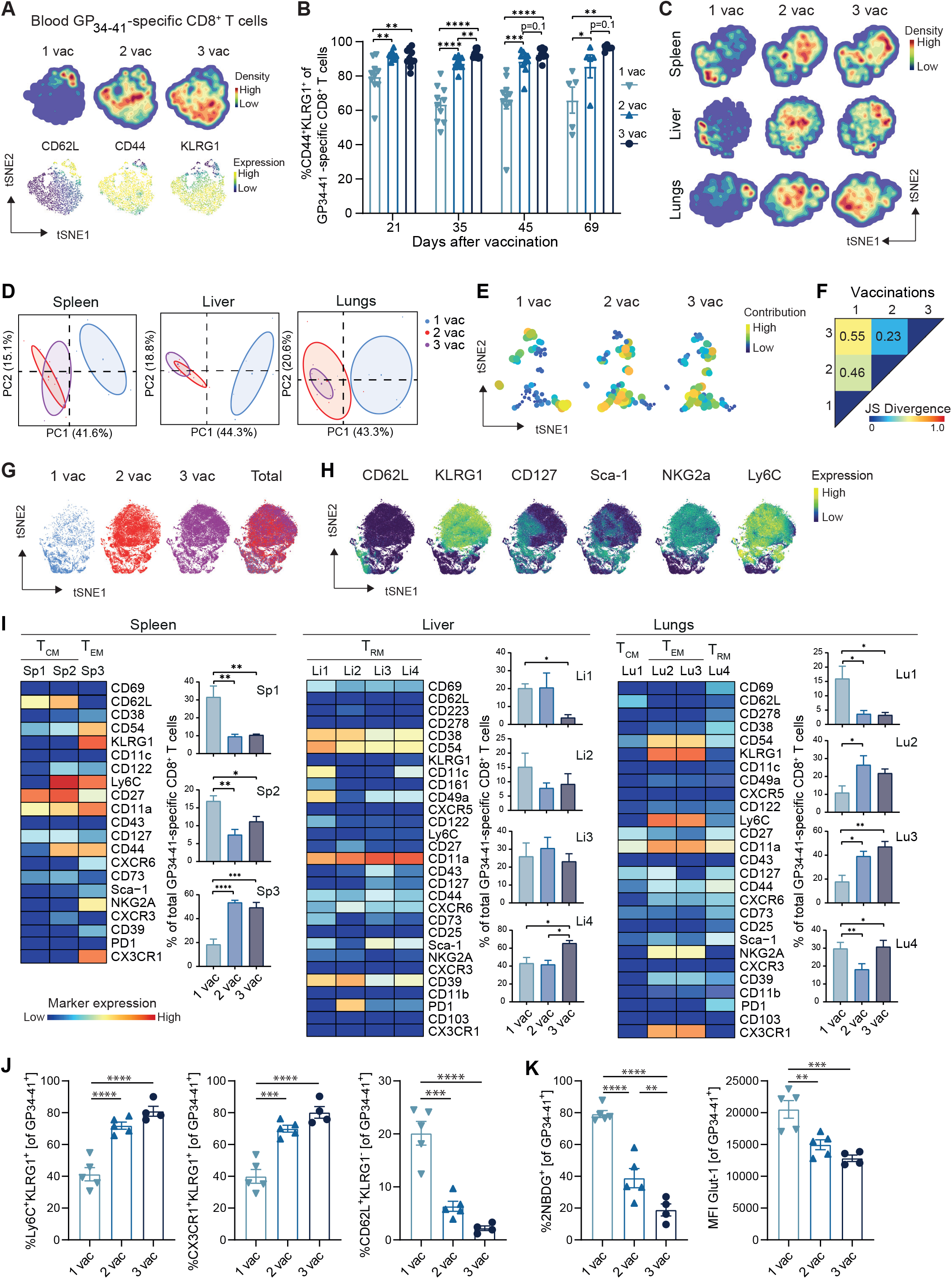
Boosting impact on the phenotype of vaccine-specific CD8^+^ T cells. **(A)** tSNE maps describing the local probability density of GP_34-41_-specific CD8^+^ T cells stained for CD62L, CD44, KLRG1 at day 71 after 1, 2 and 3 vaccinations. **(B)** Frequencies of CD44^+^KLRG1^+^ GP_34-41_-specific CD8^+^ T cells in blood. Bar graphs represent means ± SEM. Symbols represent individual mice. **(C)** tSNE maps describing the local probability density of GP_34-41_-specific CD8^+^ T cells in spleen, liver and lungs stained at day 71 after 1, 2 and 3 vaccinations with the mass cytometry panel. **(D)** Principal Component Analysis illustrating the phenotypic dissimilarity of GP_34-41_-specific CD8^+^ T cells per tissue upon multiple vaccinations. **(E)** tSNE maps showing GP_34-41_-specific CD8^+^ T cell clusters per vaccination. Clusters with similar composition profiles across samples end up close together in the map. The varying dot size and color in this cluster tSNE map shows the average cluster normalized frequencies per vaccination/tissue group. **(F)** Pairwise Jensen-Shannon Divergence plots of the tSNE map obtained from all samples of GP_34-41_-specific CD8^+^ T cells grouped by vaccinations. **(G)** tSNE embedding of GP_34-41_-specific CD8^+^ T cells isolated from vaccinated mice and multiple tissues in one analysis. Cells are color coded per vaccination. **(H)** Expression intensity of the cell-surface markers on the GP_34-41_-specific CD8^+^ T cells. The color of the cells indicates ArcSinh5-transformed expression values for a given marker analyzed. **(I)** Heat maps of GP_34-4Ĩ_-specific CD8^+^ T cell clusters in the spleen, liver and lungs of mice that received multiple vaccinations. Clusters were selected based on their significant difference and categorized in T_CM_, T_EM_ and T_RM_ subsets. Bar graphs indicate the average percentage (+ SEM) of each CD8^+^ T cell cluster within the GP_34-41_-specific CD8^+^ T-cell population elicited by one, two and three vaccinations. **(J)** Cell surface marker (Ly6C^+^KLRG1^+^, CX3CR1^+^KLRG1^+^and CD62L^+^KLRG1^-^) expression on GP_34-41_-specific CD8^+^ T cells at day 66 in spleen after 1, 2 or 3 vaccinations. Bar graphs represent means ± SEM. Symbols represent individual mice. Representative data of two independent experiments. **(K)** Glucose analog 2-(N-(7-nitrobenz-2-oxa-1,3-diazol-4-yl)amino)- 2-deoxyglucose (2-NBDG) uptake and GLUT1 expression of splenic GP_34-41_-specific CD8^+^ T cells at day 66 in after 1, 2 and 3 vaccinations. Symbols represent individual mice. Bar graphs represent means ± SEM. Representative data of two independent experiments. One-way ANOVA test with repeated measurements was used for statistical analysis. *P<0.05, **P<0.01, ***P<0.001, ****P<0.0001.

To gain further insight into the phenotype of the vaccine-specific CD8^+^ T cells upon sequential vaccination, we studied these cells in the tissues using single-cell high dimensional mass cytometry with a 39-antibody panel targeting markers of cellular activation and differentiation (31) (**Supplementary Fig 2C, Supplementary Table 1**). Visualization of the distribution of GP_34-41_-specific CD8^+^ T cells in one tSNE analysis based on density and marker expression indicated a differential distribution of these cells upon booster vaccination in the spleen, liver and lungs. While the 1^st^ vaccination was phenotypically disparate from the 2^nd^ vaccination and 3^rd^ vaccination, the 2^nd^ and 3^rd^ vaccination were overlapping but still showed some distinction (**Fig 4C**). To capture this pattern, principal component analysis (PCA) was performed based on the cluster frequencies of the GP_34-41_-specific memory CD8^+^ T cells in the spleen, liver and lungs and this confirmed the clear distinction between clusters present in one-time vaccinated *versus* clusters in two or three times vaccinated mice (**Fig. 4D**). Subsequently, we performed a dual tSNE analysis on all samples of GP_34-41_-specific CD8^+^ T cells and visualized the segregation based on vaccination-associated patterns in more depth (**Fig. 4E**). Visualization of the tSNE map values of the GP_34-41_-specific CD8^+^ T cell clusters corroborated that the 2^nd^ and 3^rd^ vaccination-specific phenotypes had considerable overlaps. To establish the (dis)similarity of the CD8^+^ T cell clusters in multiple vaccinations a Jensen-Shannon (JS) divergence analysis was performed (**Fig 4F**). A low JS distance was found when comparing antigen-specific CD8^+^ T cells induced upon 2^nd^ and 3^rd^ vaccinations, indicating a high similarity between cells induced by these vaccinations, whereas a higher JS distance was found when comparing 1^st^ to 2^nd^ vaccinations or 1^st^ to 3^rd^ vaccinations.

Visualization of the particular marker expression of the total GP_34-41_-specific CD8^+^ T cell population in the tissues indicated that the activation-associated NK cell receptors KLRG1 and NKG2A, the Ly6 family GPI-anchored surface molecules Sca-1 and Ly6C, the chemokine receptor CX3CR1 and the ectonucleotidase CD39 connected more to the 2^nd^ and 3^rd^ vaccination while CD62L expression related to single-dose vaccination (**Fig 4G, 4H**). To gain further insight into the antigen-specific CD8^+^ T cell phenotypes in the different organs, we analyzed the phenotype of these cells per tissue using FlowSOM (32) and Cytofast (33, 34), and categorized the large clusters into the three main memory T cell subsets (i.e. T_CM_, T_EM_, and T_RM_) based on their CD69 and CD62L expression. In the spleen and lungs, the percentage CD69^-^CD62L^-^ T_EM_-like cells expressing KLRG1, NKG2A, Sca-1, Ly6C, CX3CR1 within the GP_34-41_-specific CD8^+^ T cell population, increased primarily upon the first booster vaccination, while the CD69^-^CD62L^+^ T_CM_ subsets decreased (**Fig 4I**). The phenotype of CD69^+^ GP_34-41_-specific CD8^+^ T_RM_ cells in the liver was slightly affected upon the 2^nd^ boosting, and resulted in reduction of CD11c^+^CD122^+^PD-1^-^ T_RM_ cells and an increase in CD11c^+^CD122^-^PD-1^+^ T_RM_ cells (**Fig 4I**).

To validate the strong induction of the T_EM_-like phenotype by booster vaccination, we determined the percentage of cells expressing the T_EM_ markers, which provided the most distinction (i.e. KLRG1, CX3CR1, Ly6C), by flow cytometry. Indeed, vaccine-specific CD8^+^ T cells in the spleen co-expressing Ly6C^+^KLRG1^+^ and CX3CR1^+^KLRG1^+^ increased most profoundly upon the first booster and this further increased upon the second booster while the CD62L^+^KLRG1^-^ cells followed an opposite pattern **(Fig 4J)**. Moreover, this effect was also apparent upon vaccination with the Spike_539-546_-SLPs (**Supplementary Fig 2D**). Finally, the metabolic effector-memory cell properties were analyzed. Uptake of the glucose analog 2-(N-(7-nitrobenz-2-oxa-1,3-diazol-4-yl)amino)-2-deoxyglucose (2-NBDG) and expression of the glucose transport (GLUT-1) (**Fig 4K**) reduced following each vaccination, indicating lower glycolysis, which may relate to a different metabolic efficiency and to a non-exhausted state (35). Thus, the second vaccination had a profound system-wide impact on the phenotype of the vaccine-induced CD8^+^ T cells, which resulted in the increment of effector-memory-like cells characterized by activation markers and chemokine receptors.

### A third vaccination triggers the remigration of T_RM_ cells into the circulation

To fate map the T_RM_ cell progeny upon booster vaccination, we used a reporter system based on the T_RM_-restricted transcription factor Hobit to evaluate the development of the T_RM_ cells (36). Here, Hobit lineage tracer (LT) mice, which were generated by crossing CD45.1 Hobit reporter OT-I mice with ROSA26-eYFP mice, were used. CD8^+^ T cells in these mice recognize the epitope SIINFEKL (OVA_257-264_) and report active Hobit expression (by tdTomato expression) and previous Hobit expression (by YFP expression), which allows the detection of ex-Hobit^+^ cells (ex-T_RM_; tdTomato^-^ YFP^+^). First, we tested vaccination with OVA_257-264_-SLP in a prime-boost-boost setting, and this resulted in vaccine-specific CD8^+^ T cell responses that are comparable to other SLPs (**Fig 5A**). Next, we adoptively transferred the OT-I CD8^+^ T cells of the Hobit reporter × ROSA26-eYFP LT mice into naïve wild-type mice that received subsequent OVA_257-264_-SLP vaccinations in a prime-boost-boost setting. As expected, an increase of CD8^+^ T_RM_ cells (CD69^+^ tdTomato^+^ YFP^+^) was observed in the liver after the 2^nd^ and 3^th^ vaccination, which were all co-expressing CD38 (**Fig 5B, 5C)**. Remarkably, in particular the 3^rd^ vaccination induced the emergence of ex-TRM cells in the liver, spleen and blood circulation, and the formation of these cells was characterized by a T_EM_ phenotype (CD44^+^CD62L^-^KLRG1^+^) (**Fig 5C-F**). Thus, ex-TRM cells with a T_EM_ phenotype are induced upon sequential SLP vaccination and are present in the blood circulation, spleen and liver.

**Figure 5.**
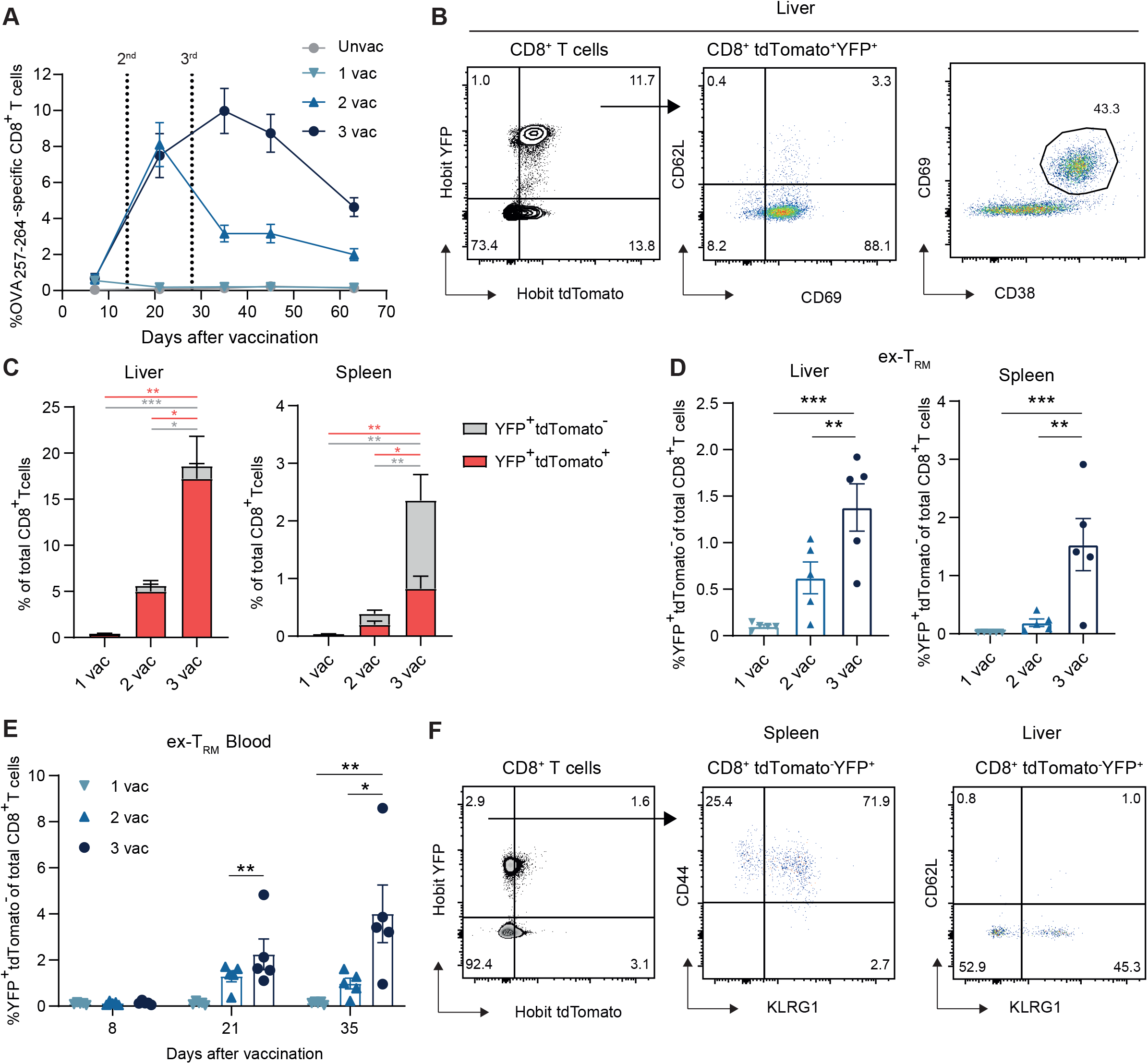
A third vaccination triggers the remigration of T_RM_ cells into the circulation. **(A**) C57BL/6 mice were vaccinated with the OVA_257-264_ T cell SLP vaccine adjuvanted with CpG in a primeboost-boost regimen with 2 week intervals. Shown are the OVA_257-264_-specific CD8^+^ T cell kinetics in blood at indicated days after vaccination. Data is represented as mean ± SEM. **(B-F)** 1 × 10^4^ OT-I Hobit LT CD8^+^ T cells were adoptively transferred into C57BL/6 mice. The day after, mice received the first OVA_257-264_-SLP vaccination, followed by 2 boosters. After 50 days spleen and liver were analyzed by flow cytometry. **(B)** Flow cytometry plots showing the CD62L, CD69 and CD38 expression of liver YFP^+^ tdTomato^+^ CD8^+^ T cells. **(C)** Percentage of CD8^+^ T cells positive YFP and/or tdTomato in liver and spleen. Data are represented as mean + SEM. One-way ANOVA was used for statistical analysis. **(D)** Percentage of tdTomato^+^YFP^+^ (ex-T_R_M) of total CD8^+^ T cells in liver and spleen. Bar graphs represent means ± SEM. Symbols represent individual mice. **(E)** Percentage of Hobit tdTomato^+^YFP^+^ (ex-T_R_M) cells of total CD8^+^ T cells in blood. Bar graphs represent means ± SEM. Symbols represent individual mice. **(F)** Phenotype of Tomato^-^YFP^-^ CD8^+^ T cells and ex-T_R_M CD8^+^ T cells in spleen and liver. One-way ANOVA was used for statistical analysis. *P<0.05, **P<0.01, ***P<0.001.

## Discussion

Here, we have investigated the capacity of single B cell and T cell epitope-containing peptide vaccines to elicit protection against SARS-CoV-2 infection in the K18-hACE2 transgenic mouse model, and found that only a third vaccination with a long peptide harboring a single T cell epitope provided full protection. To our knowledge this is the first study to show that vaccine-elicited CD8^+^ T-cells without the aid of virus-specific CD4^+^ helper T cells or neutralizing antibodies can protect against SARS-CoV-2, albeit provided in a booster setting requiring at least 2 boosters. These results may be of interest for the current discussion regarding a third vaccination, as the current vaccines elicit virus-specific T cells (4). In addition, these results may also guide the development of T-cell focused vaccines, which could be of utmost importance for susceptible patient groups such as transplantation and leukemia patients, and patients with autoimmune diseases treated with anti-CD20 antibodies such as rituximab. These patients have impaired antibody responses, and therefore may depend on robust T cell-eliciting vaccines for protection.

The DNA vaccine platform we used encoding Spike protein was highly efficient in generating neutralizing antibodies and resulted in protection against SARS-CoV-2 infection. Although, our results indicated that antibody responses to single linear B cell epitopes are inferior for induction of neutralizing antibodies, we cannot exclude the possibility that single linear B cell epitopes exist that can elicit neutralizing antibodies. In addition, in settings in which antibodies mediate protection by other mechanisms, such as antibody-dependent cellular cytotoxicity (ADCC), antibody-dependent cellular phagocytosis (ADCP), and complement dependent cytotoxicity (CDC) (37), the B cell-SLP platform may be valuable. In this respect, the addition of CpG and IFA as adjuvants and the insertion of a CD4^+^ T helper cell epitope to the linear B cell epitope vaccine was indispensable to elicit antibody responses.

In-depth studies using the T cell epitope SLP vaccines indicated that a third vaccination resulted in superior generation of CD8^+^ T_EM_ cells in the circulation and also of CD8^+^ T_RM_ cells in liver and lungs. The CD8^+^ T cells elicited upon a third dose vaccination are functionally and phenotypically modulated as evidenced by activation-associated cell-surface markers and their polyfunctional cytokine production. The latter is in line with a recent study in humans, which indicated that a third vaccination in kidney transplant recipients leads to increased circulating polyfunctional CD4^+^ T cells (38). It would be of interest to decipher whether a third dose of the mRNA vaccine in healthy individuals, which is more effective against severe COVID-19-related outcomes compared to two doses (39), also associates to increased T cell immunity.

The increase of T_EM_ and T_RM_ cells in the lungs after the third vaccination as reported here may be critical as the lungs are the primary entry point for SARS-CoV-2. Nevertheless, the circulating vaccine-induced T cells in the spleen and blood may also contribute to protection as well as these cells can rapidly migrate into the infected lung tissue and exert effector function. Moreover, also the efficient formation of the T_RM_ cells in the liver, which was mainly observed after the third vaccination, may contribute to protection as these cells have superior potential to differentiate into ex-TRM cells (40).

In conclusion, a third vaccination with a synthetic vaccine containing a single CD8^+^ T cell epitope results in protection against SARS-CoV-2. This protection can be explained by an improved quantitative and qualitative CD8^+^ T cell response after the third vaccination that is highlighted by higher numbers of virus-specific T_EM_ and T_RM_ cells with polyfunctional cytokine capacity.

## Methods

### Mice

Wild-type C57BL/6 mice were obtained from Charles River Laboratories, Jackson Laboratory or Janvier Labs. The K18-hACE2 transgenic mice, expressing the human ACE2 receptor (hACE2) under control of the cytokeratin 18 (K18) promoter (41), were obtained from the Jackson Laboratory (B6.Cg-Tg(K18-ACE2)2Prlmn/J), and bred in-house. The Hobit reporter × ROSA26-eYFP LT mice were previously described (36). At the start of the experiments, male and female mice were 6-8 weeks old. Animals were housed in individually ventilated cages under specific-pathogen free conditions at the animal facility at the Leiden University Medical Center (LUMC). All animal experiments were approved by the Animal Experiments Committee of LUMC and performed according to the recommendations and guidelines set by LUMC and by the Dutch Experiments on Animals Act.

### Peptide and DNA-based vaccination

SARS-CoV-2 linear B cell epitopes, Spike_539-546_-SLP, GP_34-41_-SLP and OVA_257-264_-SLP (aa sequences in Supplementary Table 2) were produced at the peptide facility of the LUMC. The purity of the synthesized peptide (75–90%) was determined by HPLC and the molecular weight by mass spectrometry. For the linear B cell epitope studies, mice were vaccinated subcutaneously (s.c.) at the tail base with 150 μg SLP + 20 μg CpG (ODN 1826, InvivoGen) dissolved in PBS and emulsified at a 1:1 ratio with Incomplete Freunds Adjuvant (Sigma Aldrich). For the CD8^+^ T cell SLPs, mice were vaccinated s.c. at the tail base with 100 μg of the Spike_539-546_, GP_34-41_ or OVA_257-264_-SLP + 20 μg CpG (ODN 1826, InvivoGen) dissolved in PBS. Booster vaccinations were provided with 2 weeks interval. For DNA-based vaccination, a codon-optimized, synthetic gene encoding the Spike protein of SARS-CoV-2 was generated. Plasmids were propagated in *E. coli* cultures and purified using Nucleobond Xtra maxi EF columns (Macherey-Nagel) according to manufacturer’s instructions. For vaccination, plasmids were column-purified twice, each time using a fresh column. Mice were intradermally vaccinated at the tail base with a total volume of 30 μL, containing 50 μg DNA in Tris-buffered saline (1 mM Tris, 0.9% NaCl). Booster vaccinations were provided with 3 weeks interval.

### SARS-CoV-2 infection

Clinical isolate SARS-CoV-2/human/NLD/Leiden-0008/2020 (here named SARS-CoV-2) was obtained from a nasopharyngeal sample. The next-generation sequencing data of this virus isolate is available under GenBank accession number MT705206.1 and shows one mutation in the Leiden-0008 virus Spike protein compared to the Wuhan spike protein resulting in Asp>Gly substitution at position 614 (D614G). In addition, several non-silent (C12846U and C18928U) and silent mutations (C241U, C3037U, and C1448U) in other genes were found. Isolate Leiden-0008 was propagated and titrated in Vero-E6 cells. K18-hACE2 transgenic mice were anaesthetized with isoflurane gas and intranasally infected with 5 × 10^3^ plaque forming units (PFU) of SARS-CoV-2 in a total volume of 50 μl DMEM. Mouse weight and clinical discomfort were monitored daily. All experiments with SARS-CoV-2 were performed in the Biosafety Level 3 (BSL3) Laboratories at the Leiden University Medical Center.

### Antigen-binding ELISA

ELISAs were performed to determine antibody titers in sera. Nunc ELISA plates were coated with 1 μg/ml Spike S1+S2ECD-His recombinant protein (SinoBiologicals) in ELISA coating buffer (Biolegend) overnight at 4°C. Plates were washed five times and blocked with 1% bovine serum albumin (BSA) in PBS with 0.05% Tween-20 for 1h at room temperature. Plates were washed and incubated with serial dilutions of mouse sera and incubated for 1h at room temperature. Plates were again washed and then incubated with 1:4000 dilution of horse radish peroxidase (HRP) conjugated antimouse IgG secondary antibody (SouthernBiotech, cat. 1030-05) and incubated for 1h at RT. To develop the plates, 50 μL of TMB 3,3=,5,5=tetramethylbenzidine) (Sigma-Aldrich) was added to each well and incubated for 5 minutes at room temperature. The reaction was stopped by the addition of 50 μL 1M H_2_SO_4_, and within 5 minutes the plates were measured with a microplate reader (model 680; Bio-Rad) at 450 nm

### In vitro serum neutralization titration (SNT) assay

Serum was heat-inactivated for 0.5h at 56 °C, two-fold serially diluted and incubated with 100 PFU/100μl of SARS-CoV-2 Zagreb isolate (hCoV-19/Croatia/ZG-297-20/2020, GISAID database ID: EPI_ISL_451934) for 1h at room temperature. Thereupon, 2 × 10^4^ Vero-E6 (ATCC CRL-1586) cells that were seeded in 96-well plates were inoculated with serum and virus. Following 1h incubation at 37 °C and 5% CO_2_, the inoculum was removed, and 1.5% methylcellulose overlay was added. Infected cells were incubated for 3 days at 37 °C and 5% CO_2_, prior to crystal violet staining and plaque counting.

### Single-cell preparations

Peripheral blood was collected from the tail vein. Splenocytes were obtained by mincing the tissue through cell strainers. Blood cells and splenocytes were depleted of erythrocytes using ammonium chloride lysis buffer. To remove remaining circulating blood cells from the liver and lungs, mice were perfused with 20 ml PBS containing 2 mM EDTA. Next, liver and lungs were cut into small pieces using surgical knives, and the tissue was resuspended in 3.5 ml or 1 ml, respectively, of IMDM containing 250 U/ml collagenase type 1-A (C2674, Sigma) and 20 μg/ml DNase I (D5025, Sigma). After incubation with collagenase/DNase for 25 minutes at 37°C, liver and lung tissue was dissociated into single-cell suspensions using 70 μm cell strainers, and subsequently lymphocytes were isolated using a Percoll (GE Healthcare) gradient.

### Flow cytometry

Fluorescently labelled antibodies against the following mouse antigens were used: CD3 (clone 145-2C11, BD Biosciences), CD4 (clone RM4-5, BioLegend), CD8 (clone 53-6.7, BioLegend), KLRG1 (clone 2F1, ThermoFisher), CD44 (clone IM7, BioLegend), CD62L (clone MEL-14, BioLegend), CD69 (clone H1.2F3, BD Biosciences), CX3CR1 (clone SA011F11, Biolegend), Ly6C (clone HK1.4, Biolegend), Eomes (clone Dan11mag, Invitrogen), TNF (clone MP6-XT22, Biolegend), IFN-γ (clone XMG1.2, ThemoFisher), GLUT-1 (clone EPR3915, Abcam). Zombie Aqua (Biolegend) staining was used to exclude dead cells. Glucose uptake was measured by the uptake of glucose analogue 2-(N-(7-nitrobenz-2-oxa-1,3-diazol-4-yl)amino)-2-deoxyglucose (2-NBDG, Life Technologies). Cell surface and intracellular cytokine stainings of splenocytes and blood lymphocytes were performed as described (42). PADRE-specific CD4^+^ T cells were quantified using an MHC class II tetramer for the epitope (AKFVAAWTLKAA) (NIH Tetramer Core Facility). Spike_539-546_-specific, GP_34-41_-specific or OVA_257-264_-specific CD8^+^ T cells were quantified using MHC class I tetramers generated in-house. For examination of intracellular cytokine production, single cell suspensions were stimulated with short peptides for 5h in the presence of brefeldin A or with long peptides for 8h of which the last 6h in presence of brefeldin A (Golgiplug; BD Pharmingen). Flow cytometric acquisition was performed on a BD Fortessa flow cytometer (BD Biosciences) or Aurora Cytek spectral analyzer, and samples were analyzed using FlowJo software (TreeStar).

### Mass cytometry and analysis

Metal-conjugated antibodies were either purchased from Fluidigm or were generated by conjugation of lanthanide metal isotopes to anti-mouse antibodies using the Maxpar X8 Polymer method according to the manufacturer’s protocol (Fluidigm). Cisplatins 194 and 198 and Bismuth 209 were conjugated to anti-mouse monoclonal antibodies using protocols previously described (43, 44). All in-house conjugated antibodies were diluted to 0.5 mg/ml in antibody stabilizer supplemented with 0.05% sodium azide (Candor Biosciences). Serial dilution staining was performed on mouse lymphocytes to determine appropriate antibody dilution.

The CyTOF staining was performed as described elsewhere (31). In brief, around 3 × 10^6^ cells per sample were stained for CyTOF analysis. First, cells were stained in FACS buffer with PE and APC labelled tetramers and incubated for 30 minutes on ice. Cells were washed with Maxpar Cell Staining buffer (201068, Fluidigm) and subsequently incubated for 20 minutes with 1 μM Interchalator-Rh (201103A, Fluidigm) in staining buffer. Next, aspecific binding was prevented by incubating cells with Fc blocking solution and mouse serum for 15 minutes. Anti-PE and anti-APC were added and incubated for 45 minutes. The antibody mix was added and incubated for an additional 45 minutes. After washing the cells, samples were incubated overnight with 25nM Intercalator-Ir (201192A, Fluidigm) in Maxpar Fix and Perm Buffer (201067, Fluidigm). Cells were pelleted in staining buffer and measured within one week. Before measuring, EQ™ Four Element Calibration Beads (201078, Fluidigm) were added in a 1:10 ratio. Mass cytometry data analysis focused on the antigen-specific CD8^+^ T cells. For the selection, we set our gating strategy to live single cells, positive for CD45, and excluded reference beads. The live CD45^+^ gated files were compensated using Catalyst (45). Subsequently, MHC class I tetramer-specific CD8^+^ T cells were selected in FlowJo for subsequent analysis. Next, marker expression was ArcSinh5 transformed and subjected to dimensionality reduction analyses and cluster identification using Cytosplore (30) or FlowSOM (32). For Cytosplore analysis, samples were analysed by hierarchical stochastic neighbor embedding (HSNE) (46) based on approximated t-distributed stochastic neighbor embedding (A-tSNE) (47). The similarity between tSNE maps was quantified using the Jensen-Shannon (JS) divergence as previously described (31). The JS divergence values ranged from 0 (indicating identical distributions) to 1 (indicating disjoint distributions). The dual tSNE analysis was performed to quantify the individual samples similarity based on the clusters composition as previously described (31, 48). FlowSOM was used for the identification of vaccination and tissue-specific clusters. Using FlowSOM, 14 clusters were identified per analysis. Subsequently, Cytofast (33, 34) was used for visualization and quantification of cell clusters. T_RM_ (CD62L^-^ CD69^+^), T_CM_ (CD62L^+^ CD69^-^) and T_EM_ (CD62L^-^CD69^-^) CD8^+^ T cell clusters were selected by expression of CD62L and CD69. Visualization of heatmaps and selection of clusters based on the size of the cluster (abundance of at least >5% of total) and significance was performed as previously described (31).

### Adoptive T cell transfers

Spleens of OT-I Hobit reporter × ROSA26-eYFP LT mice were isolated and subsequently CD8^+^ T cells were isolated by negative selection using MicroBeads (130-104-075, Miltenyi Biotec) according to the manufacturer’s protocol. Splenic CD8^+^ T cells were adoptively transferred via retro-orbital injection into naïve CD45.2 mice.

### Statistical analysis

Statistical analyses were performed using Cytofast or GraphPad Prism (La Jolla, CA, Unites States). One-way ANOVA and log-rank (Mantel-Cox) survival test were used for statistical analysis. All P values were two-sided, and P < 0.05 was considered statistically significant.

## Additional Information

Spike_539-546_-SLP, GP_34-41_-

Supplementary Figure 1 shows the antibody, CD4^+^ and CD8^+^ T cell responses induced by the different SLP vaccines used in this study, and the Spike DNA vaccination results. Supplementary Figure 2 shows the magnitude of Spike_539-546_- and GP_34-41_-specific CD8^+^ T cell responses in liver and lungs, the schematic of the CyTOF mass cytometry and the phenotype of the Spike_539-546_-specific CD8^+^ T cells induced by sequential vaccinations. Supplementary Table 1 shows the antibodies and metals used for CyTOF analysis. Supplementary Table 2 shows the amino acid sequences of the SLPs containing linear B cell and T cell epitopes.

## Acknowledgments

This work was funded by Health Holland (LSH-TKI project LSHM20036) and the Dutch Research Council (NWO-TTW, grant number 15830) awarded to R. Arens; LUMC (BWplus) and the graduate program of the Dutch Research Council to E.T.I. van der Gracht; Helmholtz Association’s Initiative and Networking Fund (the Project “Virological and immunological determinants of COVID-19 pathogenesis – lessons to get prepared for future pandemics (KA1-Co-02 “COVIPA”)) to L. Cicin-Sain. The authors would like to thank Prof. Dr. Jannie Borst for critically reading the manuscript, and the Experimental Animal and Flow Cytometry facilities for their support.

## Author contributions

I.N. Pardieck, E.T.I. van der Gracht, D.M.B Veerkamp, S. van Duikeren, G. Beyrend, F. Behr, J. Rip, R. Nadafi, E. Beyranvand Nejad, T.C. van der Sluis, N. Mulling, D. Brasem, M.G.M. Camps performed the experiments and analyzed the data; I.N. Pardieck, E.T.I. van der Gracht and R. Arens designed the experiments; Y. Kim and L. Cicin-Sain performed the antibody neutralization studies; S. K. Myeni, P.J. Bredenbeek, M. Kikkert contributed to the SARS-CoV-2 challenge studies; T. Abdelaal performed bioinformatics analyses; K.L.C.M. Franken, K.P.J.M. van Gisbergen and G.C.M. Zondag provided essential reagents; C.J.M. Melief, G.C.M. Zondag, J.W. Drijfhout, F. Ossendorp contributed to study design for the vaccination studies; I.N. Pardieck, E.T.I van der Gracht and R. Arens interpreted the data, prepared the figures and wrote the manuscript; R. Arens conceived and supervised the study.

