## Supplemental Figures and Tables for "A third vaccination with a single T cell epitope protects against SARS-CoV-2 infection in the absence of neutralizing antibodies"

**Figure Legends Supplementary Figures.**

**Supplementary Figure 1. (A)** C57BL/6 mice were s.c. vaccinated on day 0 ($1^{st}$ vaccination), day 14 ($2^{nd}$ vaccination) and day 28 ($3^{rd}$ vaccination) with 150 ug of SLPs consisting of linear B cell epitopes adjuvanted with CpG and Inactivated Freund's Adjuvant. Spike S1/S2 ectodomain-specific IgG antibodies in blood were determined at day 42 after vaccination. Serum from a spike DNA vaccinated animal was used as a positive control. Results are depicted as the endpoint dilution. Data is represented as mean ± SEM. Symbols represent individual mice. **(B)** C57BL/6 mice were vaccinated as described in (A) with SLPs consisting of PADRE-coupled linear B cell epitopes (E-P) or equimolar controls of the epitope alone (E), or the epitope combined with PADRE (E+P). Spike S1/S2 ectodomain-specific IgG antibodies in blood at day 42 after vaccination. Results are depicted as the endpoint dilution. Bar graphs represent means ± SEM. Symbols represent individual mice. One-way ANOVA with repeated measurements was performed to determine statistical significance. **(C)** Spike S1/S2 ectodomain-specific IgG antibodies in blood at day 27 in unvaccinated mice and in $CD4^+$ T cell depleted and proficient mice that were vaccinated with SLPs consisting of linear B cell epitopes coupled to PADRE. Results are depicted as the endpoint dilution. Bar graph represents means ± SEM. Symbols represent individual mice. **(D)** Intracellular cytokine production (IFN-γ and TNF) of $CD8^+$ and $CD4^+$ T cells in the spleen at day 35 after linear B/PADRE SLP vaccination. Splenocytes were stimulated with the linear B cell epitope coupled to PADRE SLPs, PADRE alone or no peptide. **(E)** K18-hACE 2 mice were vaccinated intradermally on day 0 ($1^{st}$ vaccination), day 21 ($2^{nd}$ vaccination) and day 28 ($3^{rd}$ vaccination) with a DNA-based vaccine encoding the Spike protein. Spike S1/S2 domain-specific IgG antibody kinetics in blood at indicated days after spike DNA vaccination. Data is represented as mean ± SEM. **(F)** Weight loss kinetics in time after SARS-CoV-2. **(G)** Survival graph after SARS-CoV-2 challenge. Significance was determined by a log-rank test. **(H)** SARS-CoV-2 neutralizing capacity by antibodies after Spike DNA vaccination on day 55 was measured by WT VNA determining the inhibition of the cytopathic effect of SARS-CoV2 on Vero-E6 cells. **(I)** Intracellular cytokine production (IFN-γ and TNF) of $CD8^+$ and $CD4^+$ T cells in the spleen at day 70 after DNA vaccination. Splenocytes were stimulated with the S1 peptivator mix. **(J)** Intracellular cytokine production of splenic $CD8^+$ and $CD4^+$ T cells at day 35 post vaccination stimulated with $Spike_{539\text{-}546}$ SLP. **(K)** Endpoint dilutions of Spike S1/S2 ectodomain-specific IgG antibodies in blood at day 42 after either 1, 2 or 3 times vaccination with the SARS-CoV-2 $CD8^+$ T cell SLP. Data is represented as mean ± SEM. Symbols represent individual mice. Serum from a spike DNA vaccinated animal was used as a positive control.

**Supplementary Figure 2. (A)** Frequencies of $CD69^+$ and $CD69^-$ $Spike_{539\text{-}546}$-specific $CD8^+$ T cells in the live immune cell population in liver and lungs at day 70 after 1, 2 or 3 vaccinations. Bar graphs represent means ± SEM. One-way ANOVA was performed to determine statistical significance. **(B)** Frequencies of $CD69^+$ and $CD69^-$ $GP_{34\text{-}41}$-specific $CD8^+$ T cells in the live immune cell population in liver and lungs at day 70 after 1, 2 or 3 vaccinations. Bar graphs represent means ± SEM. One-way ANOVA was performed to determine statistical significance. **(C)** Schematic of the mass cytometric analysis of lymphocytes isolated from spleen, liver, and lungs. **(D)** Cell surface marker ($Ly6C^+KLRG1^+$, $CX3CR1^+KLRG1^+$and $CD62L^+KLRG1^-$) expression and percentage of the glucose analog 2-(N-(7-nitrobenz-2-oxa-1,3-diazol-4-yl)amino)-2-deoxyglucose (2-NBDG) uptake of $Spike_{539\text{-}546}$-specific $CD8^+$ T cells at day 70 in spleen after 1, 2 or 3 vaccinations.

Pardieck, van der Gracht *et al.* | **Supplementary figure 1**

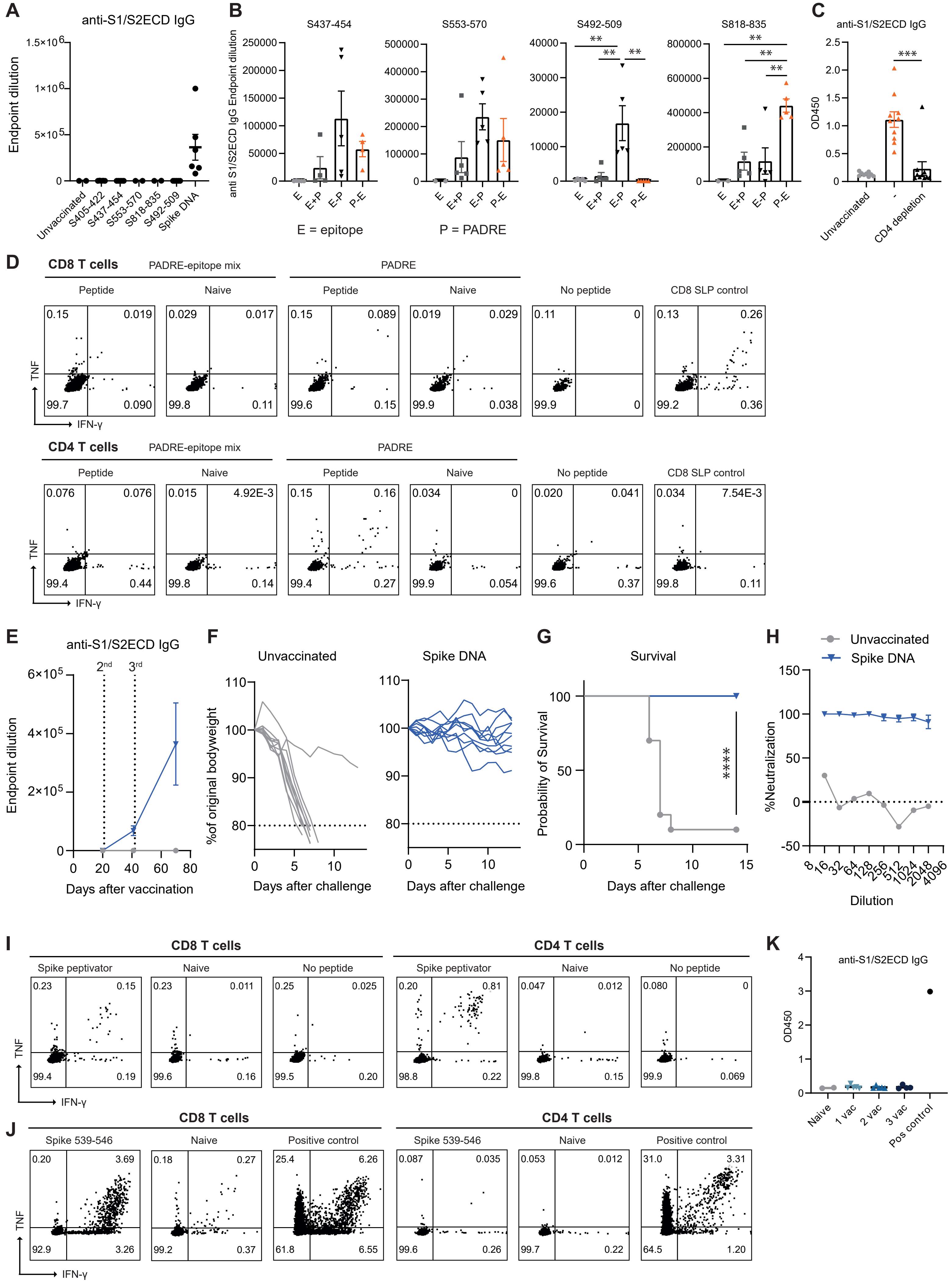

Pardieck, van der Gracht *et al.* | **Supplementary figure 2**

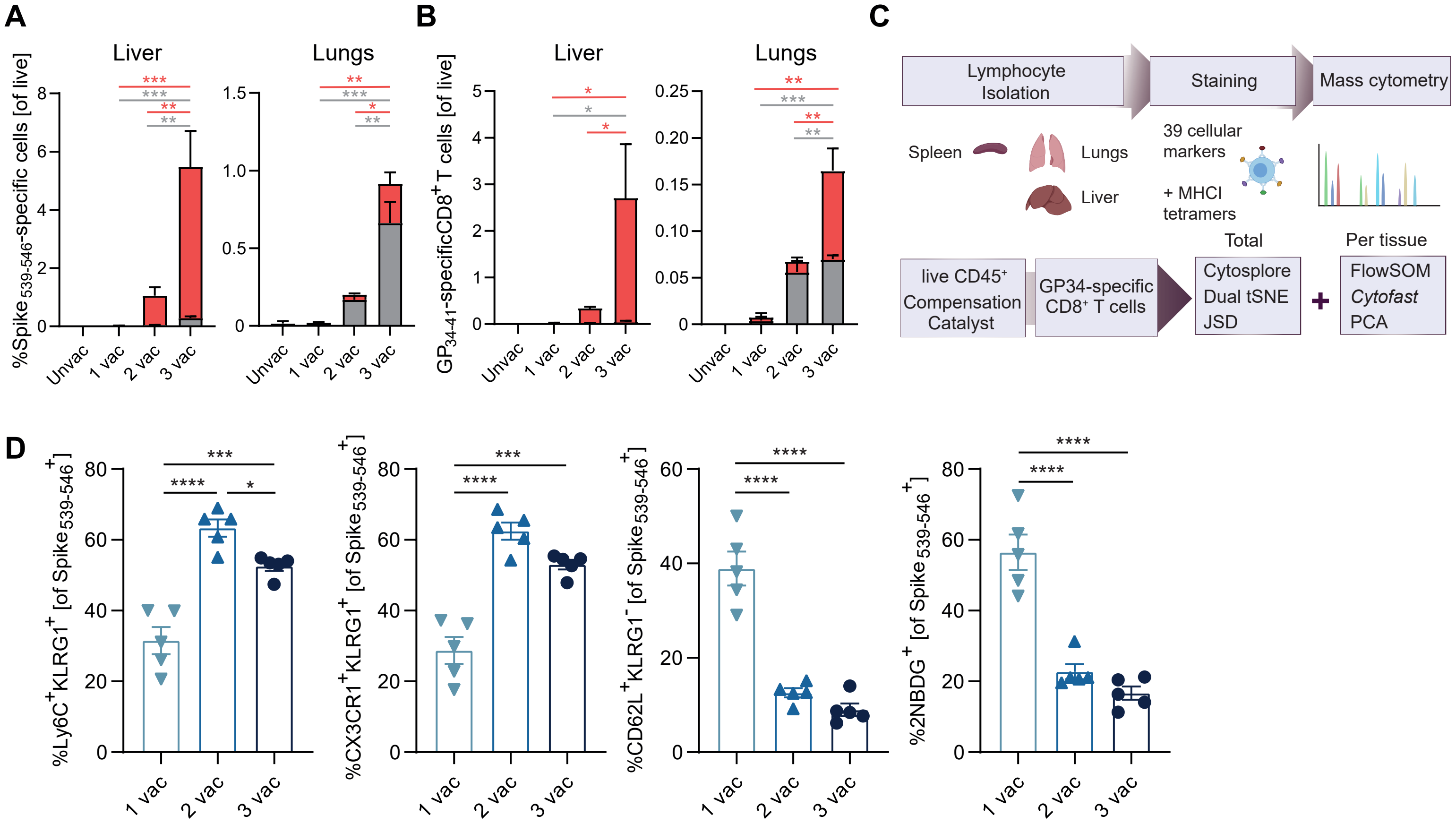

**Supplementary Table 1. CyTOF Mass Cytometry Panel**. Anti-mouse monoclonal antibodies used for staining of cells for mass cytometry analysis. Antibodies were either purchased pre-conjugated, or antibodies were conjugated to the indicated lanthanide metal isotopes

| Antibody | Clone | Metal | Pre-conjugated | Company | Cat no | Cat no metal (Fluidigm) |
|---|---|---|---|---|---|---|
| **Anti-PE** | PE001 | 165 Ho | x | Fluidigm | 3165015B | |
| **Anti-APC** | APC003 | 176 Yb | x | Fluidigm | 3176007B | |
| **CD3e** | 145-2C11 | 172 Yb | | eBioscience | 14-0031-86 | 201172A |
| **CD4** | RM4-5 | 145 Nd | x | Fluidigm | 3145002B | |
| **CD8a** | 53-6.7 | 168 Er | x | Fluidigm | 3168003B | |
| **CD8b** | YTS156.7.7 | 194 Pt | | BioLegend | 126602 | 201194 |
| **CD11a** | M17/4 | 160 Gd | | eBioscience | 16-0111-82 | 201160A |
| **CD11b** | M1/70 | 154 Sm | x | Fluidigm | 3154006B | |
| **CD11c** | N418 | 167 Er | | eBioscience | 14-0114-85 | 201167A |
| **CD19** | 6D5 | Qdot655: 112/114 Cd | x | ThermoFisher | Q10379 | |
| **CD25** | 3C7 | 150 Nd | x | Fluidigm | 3150002B | |
| **CD27** | LG.3A10 | 158 Gd | | eBioscience | 14-0272-82 | 201158A |
| **CD38** | 90 | 163 Dy | | eBioscience | 14-0381-85 | 201163A |
| **CD39** | 24DMS1 | 152 Sm | | eBioscience | 14-0391-82 | 201152A |
| **CD43** | 1B11 | 115 In | | BioLegend | 121202 | |
| **CD44** | IM7 | 142 Nd | | eBioscience | 14-0441-86 | 201142A |
| **CD45** | 30-F11 | 89Y | x | Fluidigm | 3089005B | |
| **CD49a** | Ha31/8 | 151 Eu | | BD Biosciences | 555001 | 201151A |
| **CD54** | YN1/1.7.4 | 164 Dy | | BioLegend | 116102 | 201164A |
| **CD62L** | MEL-14 | 169 Tm | | BioLegend | 104443 | 201169A |
| **CD69** | H1.2F3 | 143 Nd | x | Fluidigm | 3143004B | |
| **CD73** | TY/23 | 148 Nd | | BD Biosciences | 550738 | 201148A |
| **CD86** | GL1 | 171 Yb | | eBioscience | 14-0862-85 | 201171A |
| **CD103** | 2.E7 | 173 Yb | | eBioscience | 14-1031-85 | 201173A |
| **CD122** | TM-b1 | 155 Gd | | eBioscience | 14-1222-85 | 201155A |
| **CD127** | A7R34 | 175 Lu | x | Fluidigm | 3175006B | |
| **CD160** | 7H1 | 209 Bi | | BioLegend | 143002 | |
| **CD161** | PK136 | 170 Er | x | Fluidigm | 3170002B | |
| **CD223** | eBioC9B7W | 161 Dy | | eBioscience | 14-2231-85 | 201161A |
| **CD278** | 7E.17G9 | 162 Dy | | eBioscience | 14-9942-85 | 201162A |
| **CX3CR1** | SA011F11 | 174 Yb | | BioLegend | 149002 | 201174A |
| **CXCR3** | CXCR3-173 | 149 Sm | | eBioscience | 16-1831-85 | 201149A |
| **CXCR5** | L138D7 | 153 Eu | | BioLegend | 145502 | 201153A |
| **CXCR6** | SA051D1 | 144 Nd | | BioLegend | 151102 | 201144A |
| **FR4** | TH6 | 198 Pt | | BioLegend | 125102 | 201198 |
| **KLRG1** | 2F1 | 166 Er | | eBioscience | 16-5893-85 | 201166A |
| **Ly6C** | HK1.4 | 156 Gd | | eBioscience | 16-5932-85 | 201156A |
| **NKG2A** | 20d5 | 147 Sm | | eBioscience | 16-5896-85 | 201147A |
| **PD-1** | 29F.1A12 | 159 Tb | x | Fluidigm | 3159024B | |
| **Sca-1** | D7 | 141 Pr | | BioLegend | 108135 | 201141A |
| **TCRgd** | eBioGL3 | 146 Nd | | eBioscience | 14-5711-85 | 201146A |

**Supplementary Table 2. Peptides with Linear B cell and T cell epitopes**

X = cyclohexylalanine, u = d-Ala, B = amide

| | |
|---|---|
| PADRE | AKXVAAWTLKAuB |
| Spike$_{553-570}$ | TESNKKFLPFQQFGRDIA |
| Spike$_{553-570}$-PADRE | TESNKKFLPFQQFGRDIAAKXVAAWTLKAuB |
| PADRE-Spike$_{553-570}$ | AKXVAAWTLKAuTESNKKFLPFQQFGRDIA |
| Spike$_{818-835}$ | PSKPSKRSFIEDLLFNKV |
| Spike$_{818-835}$-PADRE | PSKPSKRSFIEDLLFNKVAKXVAAWTLKAuB |
| PADRE-Spike$_{818-835}$ | AKXVAAWTLKAuPSKPSKRSFIEDLLFNKV |
| Spike$_{405-422}$ | DEVRQIAPGQTGKIADYN |
| Spike$_{405-422}$-PADRE | DEVRQIAPGQTGKIADYNAKXVAAWTLKAuB |
| PADRE-Spike$_{405-422}$ | AKXVAAWTLKAuDEVRQIAPGQTGKIADYN |
| Spike$_{437-454}$ | NSNNLDSKVGGNYNYLYR |
| Spike$_{437-454}$-PADRE | NSNNLDSKVGGNYNYLYRAKXVAAWTLKAuB |
| PADRE-Spike$_{437-454}$ | AKXVAAWTLKAuNSNNLDSKVGGNYNYLYR |
| Spike$_{492-509}$ | LQSYGFQPTNGVGYQPYR |
| Spike$_{492-509}$-PADRE | LQSYGFQPTNGVGYQPYRAKXVAAWTLKAuB |
| PADRE-Spike$_{492-509}$ | AKXVAAWTLKAuLQSYGFQPTNGVGYQPYR |
| Spike$_{539-546}$ | IKNQCVNFNFNGLTGTGVLTESNK |
| GP$_{34-41}$ | KAVYNFATCGIFALIS |
| OVA$_{257-264}$ | SMLVLLPDEVSGLEQLESIINFEKLTEWTS |
